# Rapid imaging of lysyl oxidase activity and fibrogenesis with a turn-on fluorophore

**DOI:** 10.64898/2026.05.14.725156

**Authors:** Dongjun Li, Iván Coto Hernández, Chloe Brasket, Ibrahim Ragab Eissa, Pamela Pantazopoulos, Kenneth K. Tanabe, Jonathan C. T. Carlson, Jerrold Turner, Peter Caravan, Mariane Le Fur

**Affiliations:** Department of Radiology, Center for Interdisciplinary Innovation in Imaging, Massachusetts General Hospital, Charlestown, MA, USA; Athinoula A. Martinos Center for Biomedical Imaging, Department of Radiology, Massachusetts General Hospital, Charlestown, MA, USA; Department of Radiology, Harvard Medical School, Boston, MA, USA; Department of Surgery, Massachusetts General Hospital, Harvard Medical School, Boston, MA, USA; Center for Systems Biology and Department of Medicine, Massachusetts General Hospital, Harvard Medical School, 02114 Boston, MA, USA; Laboratory of Mucosal Pathobiology, Department of Pathology, Brigham and Women’s Hospital, Harvard Medical School, Boston, MA, USA

**Keywords:** Allysine, extracellular matrix, lysyl oxidase, fibrogenesis, histology, click chemistry, turn-on fluorescence dye

## Abstract

Fibrogenesis is essential to wound healing, but aberrant fibrogenesis is a driver of many chronic diseases and cancers. Lysyl oxidases (LOX) play a pivotal role in fibrogenesis by catalyzing the oxidation of lysine residues to reactive aldehydes (allysine) in collagens and elastin, resulting in the crosslinking and excessive deposition of these extracellular matrix components. Currently, rapid and robust histological assays to visualize the spatial distribution of LOX activity are lacking, hindering the precise validation of anti-fibrotic therapies. Here, we present a histological fluorescent staining method to visualize fibrogenesis (active fibrosis) and LOX activity in tissue sections utilizing a bioorthogonal tag and a click reaction with a turn-on fluorophore. Notably, requiring only two commercial reagents, this protocol can be completed in under two hours and is compatible with other imaging modalities, including second-harmonic generation and immunofluorescence staining. We validated this method across various healthy and fibrotic mouse and human tissue specimens.

---

Fibrogenesis is a dynamic process characterized by the production and degradation of extracellular matrix (ECM) components in response to inflammation or tissue damage. Fibrogenesis is a natural process of development and is necessary for normal wound healing, but aberrant fibrogenesis results in excessive ECM accumulation, leading to pathological fibrosis and contributing to organ dysfunction and failure [1]. Fibrosis can affect virtually any organ system, including the lungs, liver, heart, intestine, and kidneys, and is a leading cause of death worldwide [2, 3]. Fibrosis is also a key component of various types of cancer, with fibrotic stroma being deeply associated with tumorigenesis, progression, metastasis, and therapy resistance [4-7]. Therapeutic strategies targeting fibrogenic pathways have been explored as a promising approach to slow down or reverse the progression of fibrosis [2, 8, 9]. Despite the pervasiveness of fibrotic disorders, most anti-fibrotic drug candidates have failed to show efficacy in clinical trials [9]. For example, currently only three anti-fibrotic drugs (pirfenidone, nintedanib, and nerandomilast) are approved for use in patients with pulmonary fibrosis, and these drugs only mitigate disease progression, rather than reversing accumulated damage.

Collagen and elastin are the primary components of the ECM and contribute to its biomechanical properties by providing the tensile strength and viscoelasticity required for proper function [10, 11]. During fibrogenesis, activated cells, such as myofibroblasts, markedly overproduce soluble ECM components, including tropocollagen and tropoelastin, and secrete LOX enzymes involved in the cross-linking of ECM proteins (Figure 1A). LOX and its paralogs LOX-like 1-4 (LOXLn), are extracellular enzymes that catalyze the oxidative deamination of the ε-amino group of selected lysine residues on collagen monomers and tropoelastin, to form highly reactive allysine (α-aminoadipic-δ-semialdehydes) [12-15]. These allysine residues subsequently undergo spontaneous condensation with adjacent lysine/hydroxylysine and allysine/hydroxyallysine residues to form covalent cross-links in fibrillar collagens and elastin [8, 16]. LOX activity is essential for the integrity of elastic and collagen fibers in healthy tissues, including lungs, skin, and vessel walls. However, expression of LOX is markedly increased in fibrotic tissues at sites of ongoing injury [17], where it serves as the primary enzymatic driver of the structural stiffening that defines clinical pathology. Currently, rapid and robust histological assays to directly visualize the spatial distribution of LOX activity and ECM cross-linking dynamics are lacking, hindering the precise validation of anti-fibrotic therapies.

**Fig. 1.**
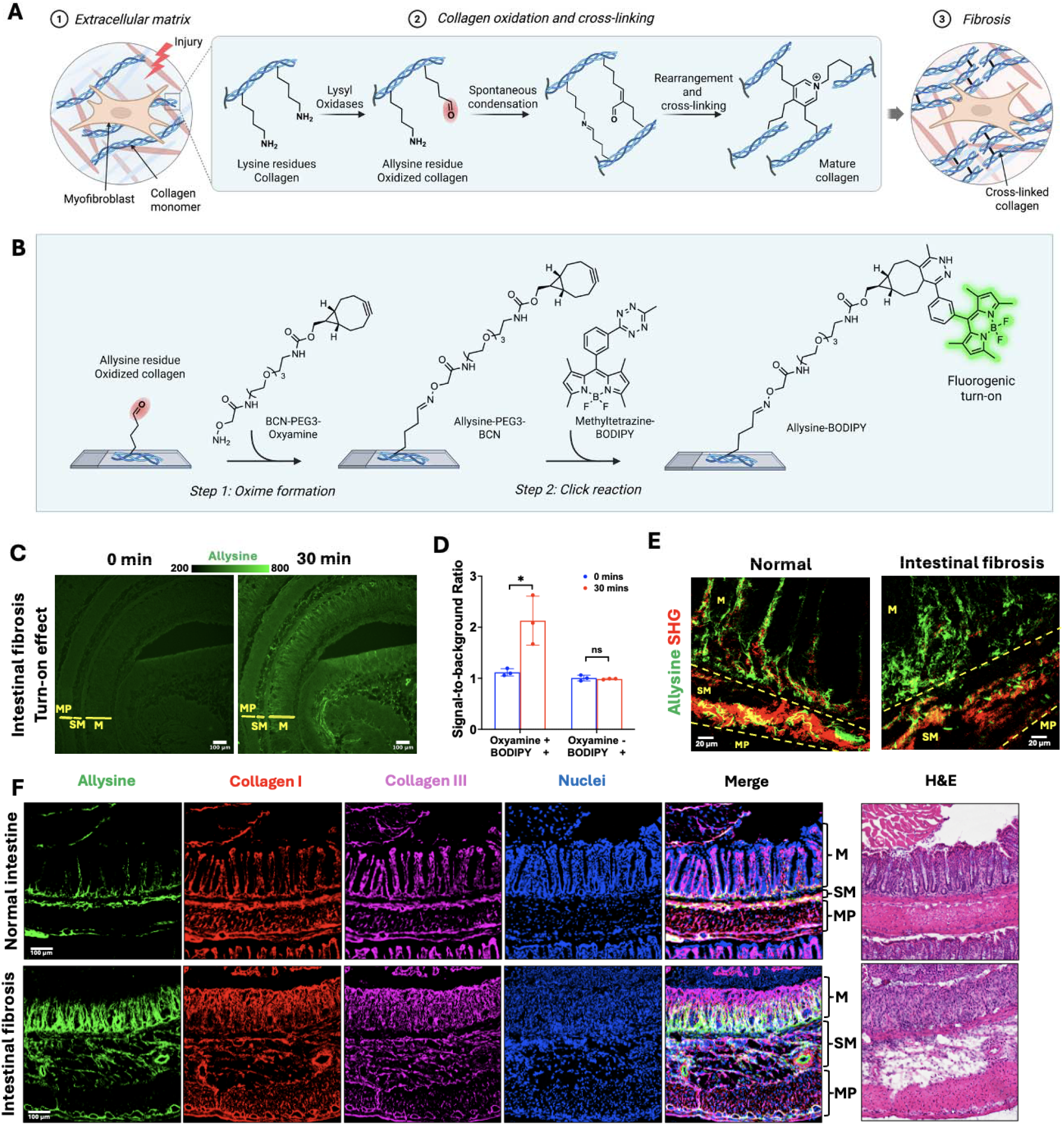
Allysine staining approach and validation. **(A)** Lysyl oxidase (LOX)-mediated oxidation and cross-linking of extracellular matrix proteins, exemplified for collagen. During fibrogenesis, LOX enzymes oxidize amino groups on lysine residues present on collagen to aldehyde to form allysine residues. Subsequently, allysine residues undergo condensation reactions with other lysine or allysine residues spontaneously to form divalent crosslinks. Further condensation of the divalent cross-link products results in tri- and tetra-cross-links, leading to mature collagen fibrils and fibers. Allysine is present only during fibrogenesis; it is absent from mature, cross-linked collagen. A similar mechanism also applies to elastin cross-linking. (Created in BioRender. LE FUR, M. (2026) https://BioRender.com/byo90fx). **(B)** Schematic illustration of the allysine staining method. First, tissue sections are incubated with an oxyamine reagent (BCN-PEG3-Oxyamine) that reacts with aldehyde groups present on allysine residues to form an oxime via a condensation reaction. The tissue sections are then rinsed in PBS to remove unreacted oxyamine reagent, followed by incubation with methyltetrazine-BODIPY. This fluorophore is almost entirely non-fluorescent at baseline but becomes exceptionally bright following the click reaction between the tetrazine and the BCN moiety. Using this activatable fluorophore enables the specific detection of allysine residues in tissue sections with minimal background signal. (Created in BioRender. LE FUR, M. (2026) https://BioRender.com/byo90fx). **(C)** The fluorogenic turn-on was demonstrated by real-time imaging in mouse intestinal sections obtained from mice with intestinal fibrosis (n = 3). Tissue sections were pre-tagged with BCN-PEG3-Oxyamine, rinsed in PBS, and then imaged immediately after the addition of methyltetrazine-BODIPY. The first image (left) was acquired right after the addition of methyltetrazine-BODIPY (within 1 min). During the 30 minutes of incubation, BODIPY was turned on following the click reaction between the methyltetrazine and BCN moieties, thus enabling the detection of allysine in tissue sections. M: mucosa; SM: submucosa. **(D)** Quantification of the allysine signal within the mucosa or submucosa relative to the tissue background signal within the muscularis propria. After 30 minutes of incubation with methyltetrazine-BODIPY, the signal-to-background ratio (SBR) increased 1.5- to 2.5-fold, depending on the local allysine concentration (n = 3). In the absence of pre-tagging BCN-PEG3-Oxyamine (Supplementary Fig. 1C), SBR of the mucosa remained unchanged after 30 minutes of incubation with methyltetrazine-BODIPY (n = 3). Data are means ± SD. \**P* < 0.05; ns, not significant; Multiple unpaired t-tests with false discovery rate adjustment. **(E)** Imaging of allysine (green) by two-photon microscopy and collagen (red) by second harmonic generation (SHG) in a single tissue section (n = 3). The allysine-BODIPY product exhibits two-photon absorption properties and can be detected in tissue sections simultaneously with mature collagen by SHG. Allysine (green) and collagen (red) distribution in the distal part of mouse colon sections is shown here. **(F)** Allysine staining and immunofluorescence staining of collagen I and collagen III were performed on the same fibrotic colon tissue section (n = 4). After imaging, the same tissue sections were stained by hematoxylin and eosin (H&E) staining. M: mucosa; SM: submucosa; MP: muscularis propria.

Existing methods to assess LOX activity in tissue sections primarily rely on immunohistochemistry (IHC) or immunofluorescence (IF). IHC staining of the LOX family is a demanding procedure requiring multiple steps and long incubation times. Moreover, it only informs on the LOX protein distribution, not on its enzymatic activity. Hydrogen peroxide, a LOX-catalyzed product, has been utilized as a biomarker for measuring LOX activity but suffers from a high endogenous background with limited specificity as a histological tool [18, 19]. Due to their diffusive, non-binding nature, LOX-activable fluorescent probes have also yet to show histological application [20]. Another strategy consists of chemically detecting allysine residues, the in situ product of LOX activity. To that end, a dinitrophenylhydrazine (DNPH) assay with IHC detection was developed and applied in lung and liver sections [21]. However, the need for signal amplification via the avidin-biotin complex makes this protocol labor-intensive with limited sensitivity. A fluorescent derivatization method coupled to liquid chromatography has been developed for measuring total allysine concentration in tissue homogenates. While this method provides accurate quantification of allysine, it does not inform on its spatial distribution [22]. Wang et al. [23] reported a biotin-conjugated hydrazide method for in situ measurement of allysine levels but had yet to validate it with standard histological methods. Aronoff et al. [24] reported a probe combining a collagen mimetic peptide and a LOX-activable variation of Pacific Blue dye to measure LOX activity. However, this probe essentially targets damaged collagen in the presence of LOX instead of measuring LOX activity and requires the synthesis of collagen peptides, thus limiting its broad applicability.

Here, we present a simple histological method to visualize the distribution of allysine in tissue sections. We reasoned that using a click chemistry approach would simplify the workflow and create a generalizable protocol that requires minimal optimization across organs, diseases, and species. Our strategy consists of first incubating the tissue section with an oxyamine-based reagent (BCN-PEG3-Oxyamine) that binds to allysine via oxime ligation, followed by a click reaction between the BCN moiety of that reagent and a tetrazine-based BODIPY activatable fluorophore (methyltetrazine-BODIPY) (Figure 1B). In its initial form, the BODIPY dye is minimally fluorescent, efficiently quenched by the methyltetrazine, but becomes brightly fluorescent upon reaction [25], therefore enabling the specific detection of allysine in tissue sections with minimal background signal. We validated our protocol against established collagen staining methods and applied it in various normal and fibrotic mouse and human tissue specimens. This protocol can help characterize active fibrosis (fibrogenesis) and distinguish it from pre-existing scar tissue. Applying it to specimens from injured and tumor-bearing organs, from both preclinical models and human patients, may facilitate the development of novel antifibrotic and antitumor therapies.

## 1. Results

### 1.1. Staining approach and method validation

To measure LOX activity and the progression of fibrogenesis, our approach consists of targeting allysine residues expressed on ECM proteins, namely collagens and elastin (Fig. 1A). The staining approach exploits click chemistry and is presented in Figure 1B. First, tissue sections are incubated for 30 minutes with an oxyamine reagent (BCN-PEG3-Oxyamine) that reacts with aldehyde groups present on allysine residues to form an oxime via oxime ligation (Figure 1B, step 1). This incubation step is performed in sodium acetate buffer at pH 4.0 to exploit the fast reaction rate between oxyamine and aldehyde groups under aqueous conditions at mildly acidic pH along with the high hydrolytic stability of the oxime product [26-28]. Then, tissue sections are rinsed in PBS to remove unreacted oxyamine reagent, followed by incubation with methyltetrazine-BODIPY in PBS for 30 minutes (Figure 1B, step 2). This fluorophore, previously reported by Carlson et al. [25], is almost entirely non-fluorescent at baseline but becomes exceptionally bright following the click reaction between the tetrazine and the BCN moiety, thus enabling the specific visualization of allysine. Moreover, this fluorogenic activation is water-compatible. Importantly, the entire procedure takes less than 2 hours to be completed.

We first validated our approach in healthy mouse intestinal sections. Allysine was observed in the mucosa and submucosa, which are intestinal layers rich in ECM components, particularly collagen [29](Supplementary Fig. 1A). When blocking all allysine residues in the tissue with a 10□mM methoxyamine solution, all allysine-positive staining was suppressed (Supplementary Fig. 1A), and quantification confirmed a complete absence of signal (Supplementary Fig. 1B).

Next, we demonstrated the specific activation of the fluorophore at sites of allysine expression using real-time imaging. Intestinal sections from dextran-sulfate sodium (DSS)-treated mice, a model of intestinal fibrosis, were used for this experiment. After pre-tagging the sections with BCN-PEG3-Oxyamine, an image was immediately acquired after the addition of methyltetrazine-BODIPY. Because this image exhibited weak autofluorescence, a long exposure time was used to highlight tissue structures (Fig. 1C). During the next 30 minutes of incubation, fluorescent signal increased in the mucosa and submucosa of the intestine, areas where collagen and elastin are present, whereas the signal in the muscularis propria remained at baseline (Supplementary movie). The signal-to-background ratio (SBR) of the BODIPY increased 1.5 – 2.5-fold, depending on the local allysine concentration and the selected imaging conditions (Fig. 1D). In contrast, without pre-tagging with BCN-PEG3-Oxyamine, BODIPY was not activated, and no signal change was observed (Fig. 1D and Supplementary Fig. 1C). These results confirm the specificity of our staining method for allysine.

### 1.2. Allysine staining protocol enables multiplexing

Next, we evaluated whether our allysine staining protocol could be used together with second harmonic generation (SHG) imaging as well as conventional immunofluorescence protocols in a single section. Cross-linked collagen can be specifically and sensitively visualized using SHG microscopy without exogenous labels thanks to collagen’s non-centrosymmetric triple-helix structure [30]. Here, we demonstrated that following fluorogenic activation, the BODIPY product exhibits two-photon absorption properties and can be simultaneously detected with cross-linked collagen by SHG microscopy, in a single section. We observed that in healthy mouse intestinal tissue sections, the allysine signal (green) was well co-localized with the SHG signal from cross-linked collagen (red) in both the submucosa and mucosa, consistent with homeostatic cross-linking (Fig. 1E, Supplementary Fig. 1D). However, in intestinal sections from DSS-treated mice, collagen fibrils appeared disrupted and scattered. Concurrently, distinct regions of intense allysine staining emerged, particularly in the mucosal layer, indicative of active tissue remodeling and collagen synthesis during fibrogenesis (Fig. 1E, Supplementary Fig. 1D).

Our protocol is also compatible with immunofluorescence (IF) staining, as demonstrated by co-staining for allysine, collagen I, and collagen III on sections of normal and fibrotic mouse colon. We performed IF staining for collagens I and III first, immediately followed by allysine staining. The distribution of allysine and cross-linked collagen was consistent with the two-photon imaging results. Co-localization of allysine and collagens was observed in healthy colon sections (Fig. 1F). In fibrotic colon sections, allysine appeared most prominent in the basal mucosa, while mature collagens I and III were primarily localized in the upper mucosa (Fig. 1F). Furthermore, allysine in the submucosa was distributed irregularly relative to collagen, sometimes appearing perpendicular to mature fibers (Supplementary Fig. 1E).

### 1.3. Allysine staining is robust and quantitative across multiple mouse organs

To investigate the versatility of our staining method, we applied it to multiple tissues from C57BL/6 naïve mice (Fig. 2A). In aorta, lung, liver and kidney sections, allysine staining revealed strong LOX activity in the aortic wall, lung alveoli, ducts and vessels. These are areas with elevated elastin concentration and little to no collagen, as demonstrated by the Verhoeff Van Gieson elastic staining and Masson’s trichrome staining, respectively (Fig. 2A). In contrast, muscle, healthy liver and kidney parenchyma do not have a prominent, dense ECM, and no allysine signal was detected in those regions (Fig. 2A). In the skin and intestine, the spatial distribution of allysine closely mirrored that of collagen (Fig. 2A). Both allysine and collagen were particularly abundant in the dermis of the skin and in the mucosal and submucosal layers of the large intestine, whereas no detectable activity was observed in the adjacent muscle layer of the skin or the muscularis propria of the intestine (Fig. 2A).

**Fig. 2.**
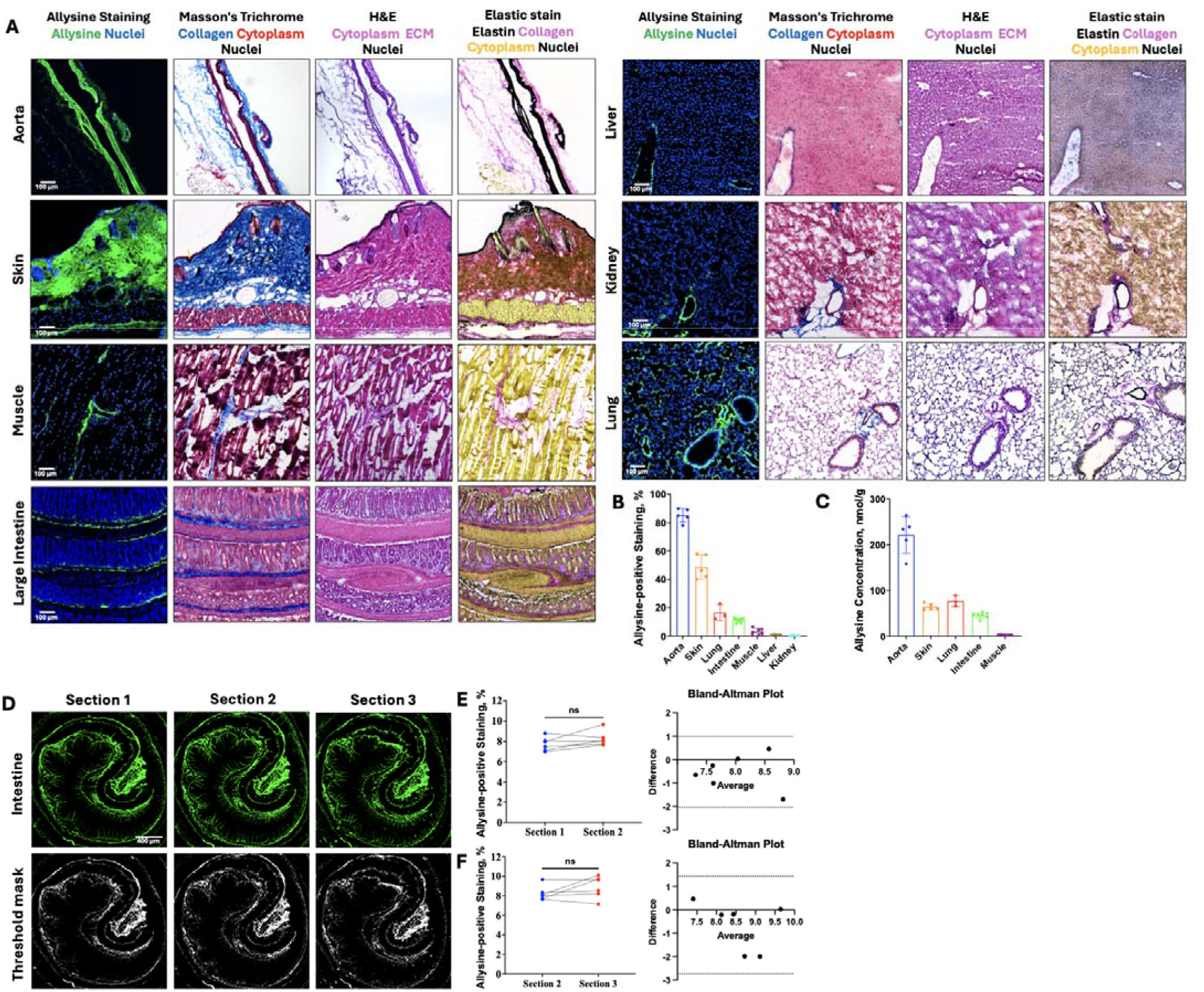
Native allysine levels across various murine organs as detected by the allysine staining. **(A)** Organs were harvested from healthy eight-week-old male C57BL/6 mice and snap-frozen for histology. Adjacent tissue sections, including aorta (n = 5), skin (n = 5), lung (n = 3), large intestine (n = 7), muscle (n = 7), liver (n = 4) and kidney (n = 4), were stained by allysine staining, Masson’s Trichrome, hematoxylin and eosin (H&E) and Verhoeff Van Gieson elastin staining. The distribution of native allysine is comparable to that of native collagen and elastin in all the organs. **(B)** Quantification of allysine-positive area percent in different organs. Data are means ± SD. **(C)** Quantification of allysine concentration in different organs using an established HPLC method. Data are means ± SD. Allysine in the liver and kidney cannot be quantified due to HPLC method limitation. **(D)** Allysine staining of three adjacent normal intestine tissue sections. Sections 1 and 2 were stained simultaneously (n = 6); Section 3 was stained one week later (n = 6). Threshold analysis highlights allysine-positive staining within the tissue. **(E)** Quantification of allysine-positive area percentage shows no significant difference between adjacent sections 1 and 2 (stained on the same day). Bland-Altman analysis indicates a bias of -0.52 between sections 1 and 2 with (SD 0.77). ns, *P* > 0.05 (paired two-tailed t test). **(F)** Quantification of allysine-positive area percentage shows no significant difference between adjacent sections 2 and 3 (stained a week apart). Bland-Altman analysis indicates a bias of -0.65 between sections 2 and 3 (SD 1.10). ns, *P* > 0.05 (paired two-tailed t test).

Next, we compared the allysine positive areas measured in sections using our methodology to the total allysine concentration measured in tissues using a biochemical assay previously reported by Waghorn et al. [22]. This assay consists of tagging allysine residues with a fluorophore during tissue digestion, followed by high-performance liquid chromatography (HPLC) for separation, detection, and quantification against a reference standard. Quantification of the percentage of allysine-positive area across entire tissue sections showed that the aorta and skin exhibited high levels of allysine (40% – 90% allysine-positive staining), while the lung and intestine showed moderate levels (10% – 20% allysine-positive staining) (Fig. 2B). In contrast, the muscle, liver, and kidney displayed little detectable allysine (0.3% – 5% allysine-positive staining) (Fig. 2B). Despite the inherent methodological differences, these results are consistent with the allysine level determined by the biochemical assay, where tissues with high molar concentration of allysine were shown to have higher allysine-positive staining using our method (Fig. 2C and Supplementary Fig. 2). Moreover, the staining could report on the spatial distribution of allysine in liver and kidney sections whereas the allysine concentrations in liver and kidney homogenates could not be determined by HPLC due to the high concentration of oxidases in these tissues which degrade allysine during the tissue digest process.

Then, to demonstrate the robustness of the allysine staining, we compared the allysine-positive area percentage obtained from adjacent serial mouse colon sections either simultaneously (sections 1 versus 2) or a week apart (sections 2 versus 3). For sections 1 and 2, quantification of the allysine-positive area percentage (IsoData algorithm, ImageJ) revealed no significant difference (P > 0.05) (Fig. 2D, E). Bland-Altman analysis confirmed an acceptable bias of -0.52 (SD 0.77) (Fig. 2E). Similarly, no significant difference was observed between the allysine-positive area percentage of sections 2 and 3 (Fig. 2D), with a Bland-Altman bias of -0.65 (SD 1.10) (Fig. 2F).

### 1.4. Allysine is a quantitative biomarker of intestinal fibrogenesis, predicting fibrosis progression

Intestinal fibrosis is a serious and common complication of inflammatory bowel disease (IBD) [9]. Currently, fibrotic strictures can only be managed by surgical resection as no anti-fibrotic drugs are clinically available for IBD patients [31, 32]. The DSS mouse model of colitis has been proposed to study intestinal fibrosis [32]. However, the natural history of fibrogenesis in this model is incompletely defined. Here, we applied our staining protocol to address this gap in knowledge.

Mice with DSS-induced colitis were sacrificed at different time points after DSS withdrawal. Masson’s trichrome staining of colon sections showed apparent collagen deposition in the mucosa and submucosa from day 11 to day 24 (Fig. 3A and Supplementary Fig. 3). Quantification of collagen-positive area in the distal colon clearly revealed that collagen deposition started on day 11 and plateaued from day 16 (Fig. 3B), consistent with prior findings [33, 34]. Analysis of adjacent sections stained for allysine revealed the upregulation of allysine in the mucosa and submucosa from day 8 to day 16 (Fig. 3A). Quantitative analysis demonstrated that fibrogenesis was initiated as early as day 8, peaked between day 11 and day 14, and resolved by day 20 (Fig. 3B) with the allysine signal returning to baseline level. The observed variation in allysine and collagen deposition aligns with the underlying mechanism of collagen oxidation and cross-linking depicted in Figure 1, and the disease progression in this model as indicated by the increase in colon weight to length ratio, a marker of disease progression (Fig.3C). As the fibrosing insult is halted (DSS withdrawal), all the allysine residues eventually become consumed as chemical cross links to form cross-linked collagen. Altogether, these data demonstrated that allysine is a quantitative biomarker of fibrogenesis; it monitors active fibrosis and predicts fibrosis progression.

**Fig. 3.**
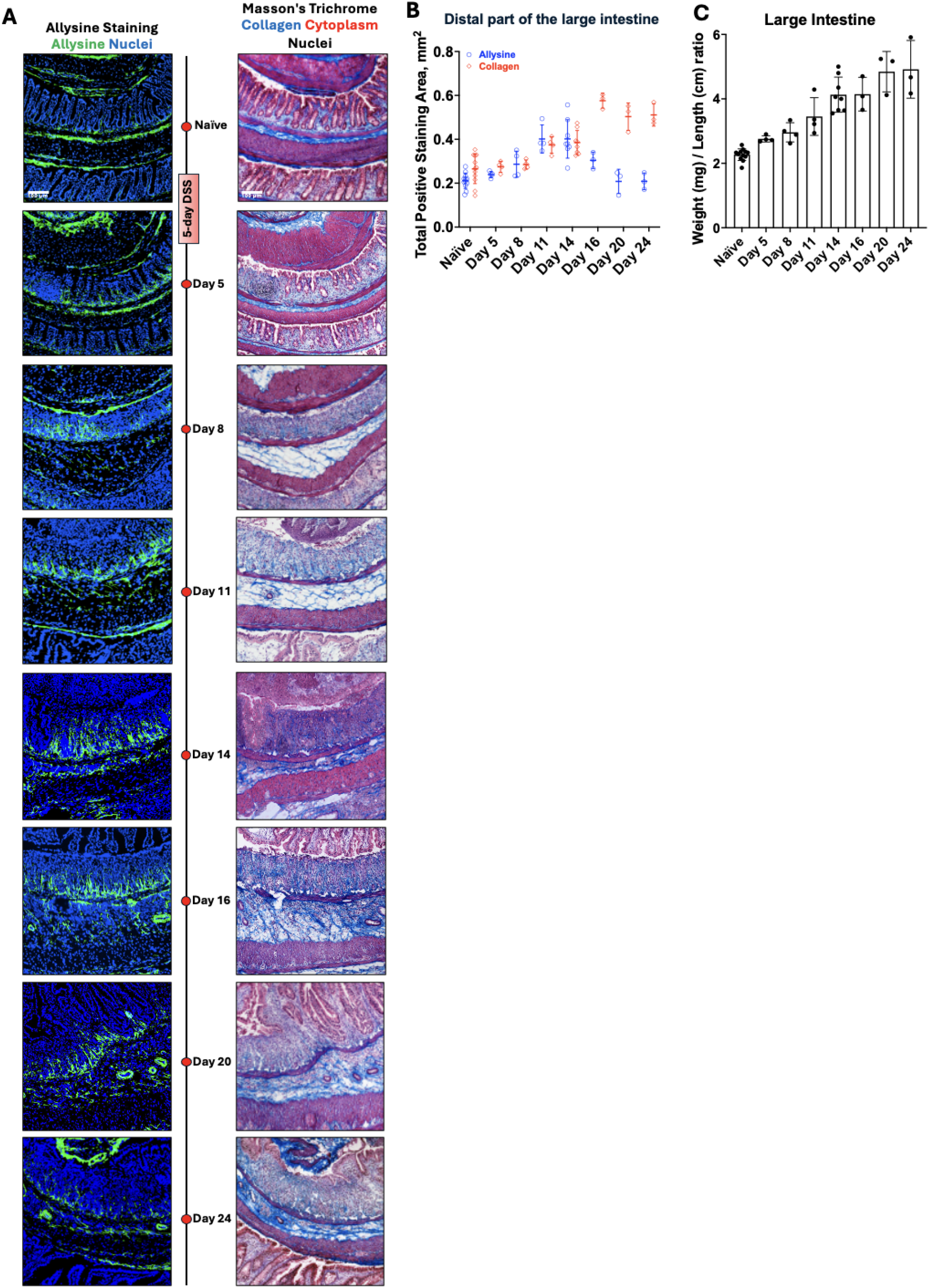
Allysine is a predictive biomarker of fibrosis. **(A)** The natural history of fibrogenesis and fibrosis in a dextran sodium sulfate (DSS) mouse model of inflammatory bowel disease. Allysine and collagen levels in the large intestine were mapped by allysine and Masson’s Trichrome staining. Eight-week-old male C57BL/6 mice received 2.5% DSS in their drinking water for five days and were subsequently sacrificed on days 5 (n = 4), 8 (n = 4), 11 (n = 4), 14 (n = 8), 16 (n = 3), 20 (n = 3) and 24 (n = 3). Age-matched naïve mice (n = 12) were used as a baseline. **(B)** Quantification of the total allysine-positive area and collagen area in the distal part of the large intestine by histological staining. Data are means ± SD. Allysine levels in the distal colon peaked on day 11-14 and decreased back to baseline level on day 20, whereas collagen levels peaked at day 16 and remained elevated. As the fibrosing insult is stopped, the allysine residues become consumed as cross-links to form cross-linked collagen. As a result, the allysine levels decrease back to baseline, whereas collagen deposition increases. This indicates that allysine is a predictor of collagen deposition. **(C)** The intestine weight-to-length ratio at different time points before and after DSS induction is an indicator of disease progression.

### 1.5. Allysine staining reports on liver fibrogenesis

Chronic liver disease, such as metabolic dysfunction-associated steatohepatitis (MASH) and alcohol-related liver disease, can lead to fibrosis and progress to cirrhosis, hepatocellular carcinoma, and ultimately, liver failure [35, 36]. Here, we applied our allysine staining method to study fibrogenesis in dietary-and toxin-induced mouse models of liver fibrosis.

We first imaged fibrogenesis in the choline-deficient, L-amino acid–defined, high-fat diet (CDAHFD) mouse model, which recapitulates key human MASH phenotypes. Male C57BL/6 mice were fed the CDAHFD for 8 weeks, at which point allysine staining revealed active fibrosis (Fig. 4A). While Sirius Red staining showed existing fibrosis in the liver, allysine staining indicated continued progression of fibrosis (Fig. 4A). In naïve control mice on a standard chow diet, the liver parenchyma was largely devoid of allysine and collagen, though both were present in the portal triad (Fig. 4A). Quantification demonstrated a 30-fold increase in allysine levels in CDAHFD livers compared to naïve controls (Fig. 4C), alongside a 10-fold elevation in collagen levels (Fig. 4D).

**Fig. 4.**
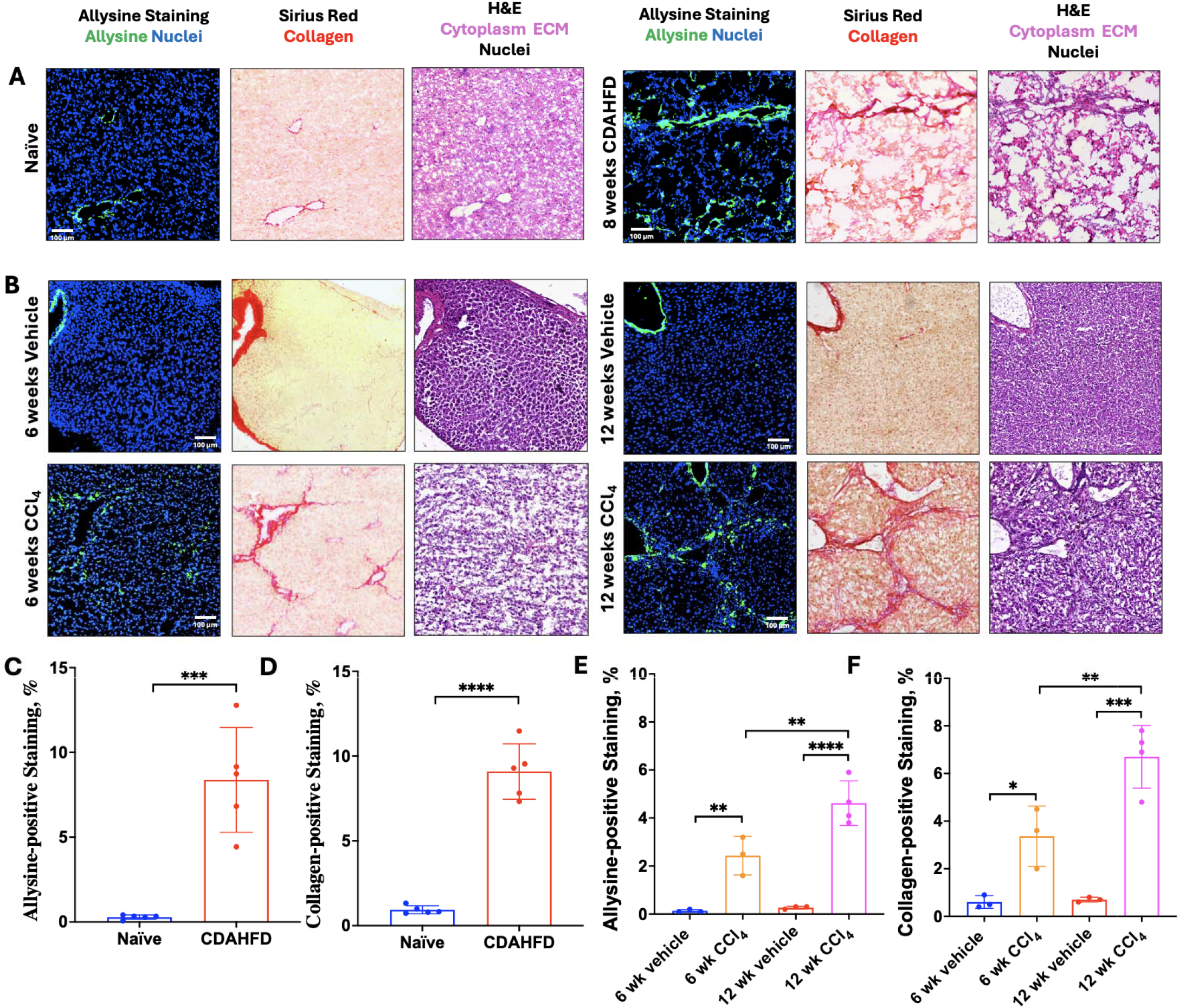
Detection of liver fibrogenesis in choline-deficient, L-amino acid–defined, high-fat diet (CDAHFD) mouse model and CCl_4_ mouse model. **(A)** Adjacent tissue sections from CDAHFD or naïve mice were stained by allysine staining, Sirius Red, and H&E staining. Allysine was present in the liver parenchyma in CDAHFD mice but not in naïve mice. **(B)** Adjacent tissue sections from CCl_4_ or age-matched vehicle mice were stained by allysine staining, Sirius Red, and H&E staining. Allysine was present in the liver parenchyma in CCl_4_ mice but not in vehicle mice. **(C)** Quantification of the allysine-positive area percent from allysine staining in CDAHFD and naïve mice (***P < 0.001, unpaired t test, two-tailed). **(D)** Quantification of the collagen-positive area percent from Sirius Red staining in CDAHFD and naïve mice (****P < 0.0001, unpaired t test, two-tailed). **(E)** Quantification of the allysine-positive area percent from allysine staining in CCl_4_ and age-matched vehicle mice. **(F)** Quantification of the collagen-positive area percent from Sirius Red staining in CCl_4_ and age-matched vehicle mice. All data are means ± SD. \**P* < 0.05; \*\**P* < 0.01; \*\*\**P* < 0.001; \*\*\*\**P* < 0.0001. one-way ANOVA followed by Tukey’s post hoc test.

To further validate our method, we examined liver tissue from mice that received either 6-weeks or 12-weeks of oral carbon tetrachloride (CCl_4_) dosing to induce liver fibrosis and compared the results to liver tissue from vehicle-treated mice using our allysine staining method. In vehicle mouse liver sections, allysine or collagen were only observed in the portal triad (Fig. 4B). After 6 weeks of CCl_4_ treatment, a mesh-like distribution of allysine was observed, and this pattern was visually similar to that seen with the collagen distribution visualized by Picrosirius Red staining (Fig. 4B). At 12 weeks, the allysine level increased approximately 1.9-fold compared to the 6-week timepoint, while the collagen level increased by approximately 2.0-fold (Fig. 4E, F). Additionally, this allysine distribution differs markedly from that observed in CDAHFD-induced liver fibrosis (Fig. 4A, B).

### 1.6. Allysine staining reveals fibrogenesis in fibrotic human liver tissue and pancreatic ductal adenocarcinoma

Next, we sought to demonstrate the applicability of mapping allysine in fibrotic human tissue specimens. First, we applied the method to visualize active fibrosis in cirrhotic and normal liver tissue. The distribution of allysine is consistent with collagen-rich fibrotic extracellular matrix in cirrhotic liver, indicating ongoing fibrosis progression (Fig. 5A and Supplementary Fig. 4). In healthy liver tissue, allysine staining maps collagen and elastin cross-linking in blood vessel walls and bile ducts (Fig. 5A and Supplementary Fig. 4).

**Fig. 5.**
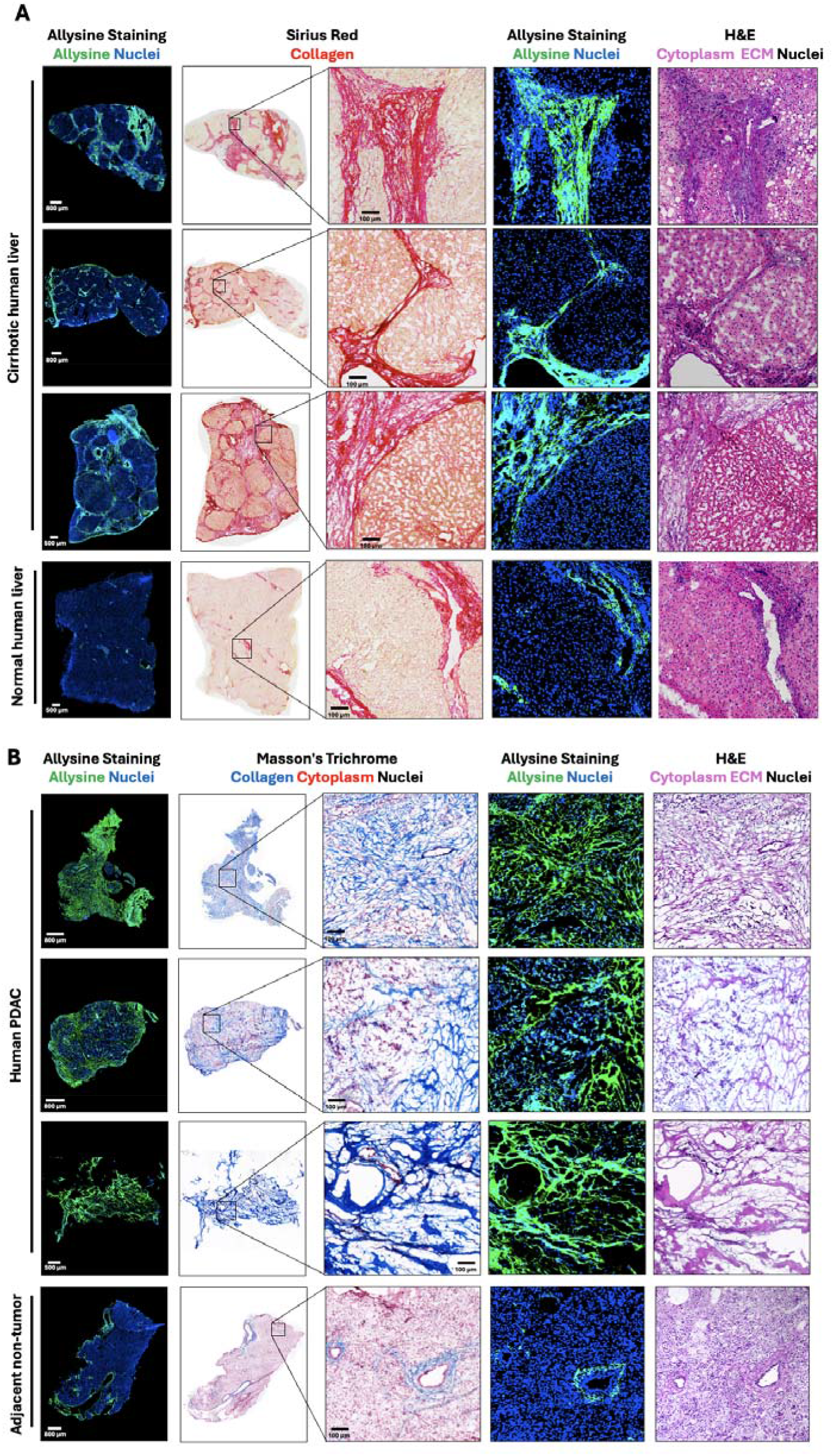
Allysine staining reveals fibrogenesis in human samples. **(A)** Cirrhotic and normal human liver specimens were obtained from patients with liver diseases. The allysine and collagen levels were mapped in the entire liver tissue section by allysine and Sirius red staining in adjacent tissue sections. Hematoxylin and eosin (H&E) staining visualized the tissue morphology. The distribution of allysine is consistent with collagen-rich fibrotic extracellular matrix. Moreover, allysine staining only maps active fibrosis in fibrotic liver tissue and maps collagen and elastin cross-linking in vessel walls and bile ducts in healthy liver tissue. **(B)** Human pancreatic ductal adenocarcinoma (PDAC) and adjacent non-tumor pancreas specimens were stained using allysine, Masson’s Trichrome and H&E stains. The distribution of allysine is consistent with the collagen-rich tumor microenvironment.

Then, we applied the allysine staining to visualize active fibrosis in human pancreatic ductal adenocarcinoma (PDAC) tissue and adjacent non-tumor pancreas. A unique feature of PDAC tumors is an excessive and dense fibrotic stroma, which influences cancer cell development and resistance to therapy. Here, fibrotic human PDAC tissue exhibited strong allysine staining, indicative of ECM remodeling (Fig. 5B and Supplementary Fig. 5) with the allysine signal co-localizing with mature collagen, as visualized by Masson’s trichrome staining. In contrast, adjacent non-tumor human pancreatic tissue was largely free of collagen and allysine, with staining confined to vessel walls (Fig. 5B and Supplementary Fig. 5).

## 2. Discussion

Our goal was to develop a robust, generalizable, and efficient staining protocol for evaluating the spatial distribution and expression of allysine residues, the in situ product of LOX activity, in tissue sections. The urgent need for such a tool arises from the lack of simple, accessible methods in understanding the mechanisms of wound healing and fibrosis progression in patients with devastating, irreversible illnesses, needed to support the development of effective anti-fibrotic therapies. Our protocol achieves this through a two-step process. First, aldehydes on allysine residues react with an oxyamine-based reagent (BCN-PEG3-oxyamine) to form an oxime via a condensation reaction. Second, a subsequent click reaction using methyltetrazine-BODIPY generates a fluorescence signal following probe activation. This two-step strategy, coupled with the solubility, stability, and reactivity of the reagents in aqueous conditions, facilitates the visualization of allysine in tissue sections.

The key advantages of this protocol are its accessibility and versatility. We purposefully selected commercially available reagents, eliminating the need for synthetic chemistry expertise and enabling widespread adoption across laboratories. The use of a superbright fluorogenic dye allows for direct visualization of allysine residues without signal amplification, thereby simplifying the process. The allysine staining protocol can be completed in under two hours, and thus, offers a significantly higher throughput compared to conventional IHC and IF protocols that typically require several days due to long incubation times. We also bypass the need for species-specific reagents by tagging aldehydes using click-chemistry reagents. We validated this method to ensure its specificity and reliability. A blocking experiment and real-time imaging confirmed the reaction specificity, while correlation with allysine concentration determined by HPLC in corresponding tissue specimens further supported signal specificity. The method proved robust across multiple mouse organs, including the aorta, skin, lung, intestine, muscle, liver, and kidney. Furthermore, the two-photon absorption properties of the activated fluorophore enable detection in single sections with mature collagen via SHG microscopy, a technique commonly used to study the organization of collagen fibrils [30]. The protocol is compatible with conventional IF staining, allowing for multiplexing and integration with existing workflows. Multiplexing enables analysis of multiple biomarkers on the same slide, thereby alleviating issues related to sample heterogeneity while increasing throughput. It is also particularly advantageous when clinical specimens are limited, for example, working with needle biopsies.

Previous research by the Caravan group established allysine as a biomarker for positron emission tomography (PET) imaging and molecular magnetic resonance imaging (MRI) [36-39]. These studies validated the in vivo efficacy of the target for detecting early-onset of fibrosis and monitoring treatment response. Notably, the staining method presented here offers superior spatial resolution compared to noninvasive PET and molecular MR imaging, providing deeper insight into allysine distribution within animal models of fibrogenesis and human fibrotic disorders. We subsequently applied this protocol to characterize the natural history of fibrogenesis and ECM deposition in the DSS mouse model of intestinal fibrosis. We demonstrated that allysine expression in the intestine was increased days before collagen deposition and that allysine is, therefore, a predictive biomarker of fibrosis. Similarly, in fibrotic mouse and human liver samples and PDAC samples, the allysine staining indicated the areas undergoing active ECM remodeling.

The main limitation of our protocol is that it requires frozen tissue samples because formalin, an aldehyde-based fixative, would interfere with allysine detection otherwise. If sample fixation is required, metharcarn (methanol-Carnoy’s fixative) could be used as an alternative to formalin, although further validation would be required. Secondly, our protocol cannot discriminate between allysine from collagen and allysine from elastin. However, this limitation could be overcome by developing a more selective targeting moiety. Nonetheless, as demonstrated in tissue staining of different organs, allysine signal from normal elastic lamina is easily differentiated from pathologic fibrogenesis because of the high microscopical spatial resolution.

In conclusion, we have developed and validated a rapid, accessible, and versatile histological fluorescent staining method to visualize allysine distribution in tissue sections. This tool could support both preclinical and clinical studies investigating novel anti-fibrotic drugs by providing a simple and reliable readout of LOX activity.

## 3. Methods

### 3.1. Reagent preparation

To prepare a BCN-PEG3-Oxyamine stock solution, 10 mL of ultrapure water was treated with 250 mg Hydroxylamine Wang resin (8.55117, Sigma) overnight under constant rotation. 10 mg of BCN-PEG3-Oxyamine powder (CP-6079, Conju-Probe) was suspended in 2 mL of Hydroxylamine Wang resin-treated water. Aliquots of the stock were filled with nitrogen gas, wrapped with Parafilm, and stored at -80 °C. A fresh working solution of 10 µM BCN-PEG3-Oxyamine was prepared in 20 mM acetate buffer with 150 mM NaCl at pH 4.0 from the stock. To prepare a methyltetrazine-BODIPY stock, 1 mg of methyltetrazine-BODIPY powder (CP-4018, Conju-Probe) was suspended in 400 μL DMSO, and aliquots were stored at -80 °C. A fresh working solution of BODIPY was prepared at 5 µM in PBS with 5% DMSO (dilute BODIPY in DMSO first and then add PBS).

### 3.2. Allysine staining for frozen tissue sections

Frozen tissue sections were rehydrated in 4 °C PBS for 10 minutes, and a waterproof circle was drawn around the tissue using a hydrophobic barrier PAP pen (H-4000, Vector). Tissue sections were subsequently fixed in methanol for 4 minutes. Sections were then washed in 4 °C PBS for 5 minutes to remove methanol. For blocking experiments only, slides were incubated with 250 µL of 10 mM methoxyamine hydrochloride in PBS for 30 minutes, and then the blocking solution was removed by shaking rigorously (without rinsing). Slides were incubated with a fresh solution of 300 µL of 10 µM BCN-PEG3-Oxyamine in 20 mM acetate buffer with 150 mM NaCl at pH 4.0 for 30 minutes at room temperature. Slides were washed with 4 °C PBS three times for 5 minutes each at 4 °C. Slides were incubated in the dark with 300 µL of 5 µM methyltetrazine-BODIPY reagent in PBS with 5% DMSO for 30 minutes. Slides were differentiated in the dark in 50% ethanol for 1 minute, 75% ethanol for 1 minute, 95% ethanol for 3 minutes, and 100% ethanol for 1 minute. Mounting medium with DAPI (H-1200, Vector) was added to the slides and covered with a coverslip. Slides were stored in the dark at room temperature for 18 hours and imaged the next day using a fluorescence microscope (Nikon Instruments). A broadband white-light source (Lumencor SOLA Light Engine) was used for fluorescence excitation. The fluorescence emission signal was separated from the excitation light using Semrock BrightLine filter cubes for DAPI (Ex 356/30 nm, Em 447/60 nm) and allysine (Ex 466/40 nm, Em 525/50 nm).

### 3.3. Real-time allysine imaging and signal quantification

Frozen colon sections (n = 3) from mice with intestinal fibrosis were rehydrated in 4 °C PBS for 10 minutes, and a waterproof circle was drawn around the tissue using a hydrophobic barrier PAP pen. Tissue sections were fixed in methanol for 4 minutes. Sections were washed in 4 °C PBS for 5 minutes. Slides were incubated with a fresh solution of 300 µL of 0.5 µM BCN-PEG3-Oxyamine in 20 mM acetate buffer with 150 mM NaCl at pH 4.0 for 30 minutes at room temperature. Slides were washed with 4 °C PBS three times for 5 minutes each at 4 °C. Tissue sections were imaged immediately after the addition of 5 µM methyltetrazine-BODIPY and imaged every 10 seconds for 30 minutes using a fluorescence microscope as described in section 3.2. To measure the baseline signal, the pre-incubation step with BCN-PEG3-Oxyamine was skipped.

The signal intensity of allysine-rich areas (SI allysine) was measured by manually drawing three regions of interest (ROIs) on an allysine-positive area (fibrotic area) in the submucosa or mucosa of the large intestine. The signal intensity of the background (SI background) was measured by placing three ROIs on the muscle area within the muscularis propria (no allysine). These ROIs were then copied to every frame of the time-lapse sequence. Signal-to-background ratio (SBR) was calculated at each time point using equation 1.

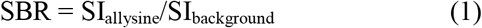

### 3.4. Multi-photon Images

A commercial multiphoton microscope (Bliq Photonics) powered by an ultrafast laser (Chameleon Vision-S, Coherent) was used. The wavelength of the ultrafast laser was tuned to 850 nm, and the beam intensity was set to 15% to simultaneously collect two-photon excitation fluorescence signal from allysine and second harmonic generation (SHG) from collagen structures. The section sample was imaged using a 20x/1.0 NA water immersion objective lens (XLUMPlanFL N 2.8 mm WD, Olympus, Japan). Multiphoton fluorescence and SHG signals were collected via non-descanned detection using a 495 nm long pass filter, a pair of band pass filters (ET520/40 and 442/42), and two photomultiplier tubes (H 7422-40, Hamamatsu) with a gain set to 80% and 95%, respectively. Images were acquired using a polygon laser scanner operating at 7.81 FPS (1024 x 512) and averaged 30 times with the same image acquisition software (Nirvana 2.9) to enhance the signal-to-noise ratio (SNR) in both imaging modalities.

### 3.5. Immunofluorescence and allysine co-staining

Frozen tissue sections were rehydrated in 4 °C PBS for 10 minutes, and a water-proof circle was drawn around the tissue using a hydrophobic barrier PAP pen. Tissue sections were subsequently fixed in methanol for 4 minutes and then washed in 4 °C PBS for 5 minutes. Tissue sections were blocked with 2% BSA (A3294, Sigma-Aldrich) in PBST (PBS with 0.1% Triton X-100) for 1 hour at room temperature. Sections were incubated with anti-collagen I antibody (1:200 dilution in blocking buffer; 1310-01, Southern Biotech) and anti-collagen III antibody (1:100 dilution; 22734-1-AP, Proteintech) overnight at 4 °C and then washed with PBST three times for 5 minutes each. Sections were incubated with donkey anti-goat secondary antibody conjugated with Alexa Fluor 555 (1:2000 dilution in blocking buffer; A-21432, Thermo Fisher) and donkey anti-rabbit secondary antibody conjugated with Alexa Fluor 647 (1:2000 dilution; A-31573, Thermo Fisher) in the dark and then washed with PBST three times for 5 minutes each. Sections were rinsed briefly in ultrapure water. Subsequently, sections were incubated in the dark with 10 µM BCN-PEG3-Oxyamine in 20 mM acetate buffer with 150 mM NaCl at pH 4.0 for 30 minutes at room temperature. Sections were washed with 4 °C PBS three times for 5 minutes each at 4 °C. Slides were incubated in the dark with 5 µM methyltetrazine-BODIPY reagent in PBS with 5% DMSO for 30 minutes. Slides were differentiated in the dark in 50% ethanol for 1 minute, 75% ethanol for 1 minute, 95% ethanol for 30 seconds, and 100% ethanol for 30 seconds. Mounting medium with DAPI was added to the slides and covered with a coverslip. Slides were imaged the next day using a fluorescence microscope (Nikon Instruments). Excitation and emission were selected using Semrock BrightLine filter cubes for DAPI (Ex 356/30 nm, Em 447/60 nm), allysine (Ex 466/40 nm, Em 525/50 nm), collagen I (Ex 554/23 nm, Em 609/54 nm), and collagen III (Ex 618/50 nm, Em 698/70 nm).

### 3.6. Tissue digestion and allysine quantification by high-performance liquid chromatography (HPLC)

Allysine was quantified in tissue samples by adapting a fluorescence derivatization protocol previously reported [22]. Briefly, tissues (50–100 mg) were digested in 1 mL of 6 M HCl with 40 mg of sodium 2-naphtol-6-sulfonate (N0035, TCI) at 105 °C overnight in a dry bath block heater. The digest was then adjusted to a pH of 2 using 6 M NaOH. pH-adjusted digest was mixed with an equal amount of DMSO for allysine quantification by HPLC. An allysine-naphtol reference standard was synthesized as previously described [22], and a 30 µM stock solution was prepared in water.

Analytical reverse-phase chromatography was performed using a Discovery® C8 (5 µm) HPLC column (59353-U, Supelco) on a 1290 Infinity II Agilent chromatography system equipped with fluorescence detection. The method was as follows: Mobile phase A: 0.1% trifluoroacetic acid (TFA) in water; mobile phase B: 0.1 % TFA in acetonitrile; flow rate: 1.6 mL/min; gradient: from 12% to 14% B in 2 min, 14% B for 8.5 min, up to 16% B in 2 min, up to 95% B in 6.5 min, 95% B for 1 min, down to 12% B in 0.5 min and 12% B for 2 min. The allysine-naphtol derivatization product was detected using λ_ex_ = 254 nm and λ_em_ = 380 nm. A 19 μL sample of interest was first injected into the column, and then standard addition was performed in the HPLC, using the Agilent injection program, by spiking the sample with a 0.2, 0.4, and 0.6 μL of 30 μM allysine-naphthol standard. The area under the curve (AUC) of the allysine-naphthol peak of each sample was obtained at ∼11.2 minutes retention time (OpenLab, Agilent) (Supplementary Fig. 4 B and C). The concentration of allysine in each sample was calculated by the standard addition method (Supplementary Fig. D and E).

#### Animal experiments

All animal experiments and procedures were performed in accordance with the National Institutes of Health Guide for the Care and Use of Laboratory Animals in compliance with the ARRIVE (Animal Research: Reporting of In Vivo Experiments) guidelines and were approved by the Institutional Animal Care and Use Committee of Massachusetts General Hospital. Frozen mouse liver samples from CDAHFD and CCl4 models of liver fibrosis, as well as corresponding naïve or sham liver samples, were provided by Dr. Caravan. These samples were banked from previous animal studies and stored to be reused for other research purposes.

### 3.7. Naïve mice

Eight-week-old male C57BL/6 mice (Charles River Laboratories) were sacrificed, and their organs (aorta, skin, muscle, large intestine, liver, kidney, lung) were harvested for histological analysis or tissue digestion for HPLC quantification.

### 3.8. Dextran sodium sulfate (DSS)-induced intestinal fibrosis mouse model

To induce colitis, eight-week-old male C57BL/6 mice (Charles River Laboratories) received 2.5% DSS (0216011090, MP Biomedicals) in autoclaved drinking water for 5 days. DSS was withdrawn, and normal autoclaved drinking water was resumed on day 6. Mice were euthanized on days 5 (n = 4), 8 (n = 4), 11 (n = 4), 14 (n = 8), 16 (n = 3), 20 (n = 3), and 24 (n = 3), and their large intestine was collected for histology. Colons were dissected from the rectum to the colonocecal margin and carefully flushed with PBS to remove intestinal content. Excess PBS was removed using gauze, and the length and weight of the colons were measured. Colons were cut longitudinally and rolled as “Swiss Rolls” from the rectum to the proximal colon, mucosa facing up.

### 3.9. Choline-deficient, L-amino acid–defined, high-fat diet (CDAHFD) mouse liver fibrosis model

Male C57BL/6 mice (8 to 10-week-old; Charles River Laboratories) received a CDAHFD (A06071302, Research Diets Inc.) containing 60% fat calories and 0.1% methionine for 8 weeks, while the naïve mice maintained standard chow. Simple randomization assigned the mice to two experimental groups: Naïve (standard diet; n = 5), CDAHFD (8-week CDAHFD; n = 5).

### 3.10. CCl_4_ liver fibrosis mouse model

Male C57BL/6 mice (6-week-old; Charles River Laboratories) were treated with an oral gavage of carbon tetrachloride (CCl_4_) for 6 weeks (n = 3) or 12 weeks (n = 4) (2–3 times per week, 0.1 mL of 20% CCl4 in olive oil the first week, 30% the second week, and 40% from weeks 3 through 6 or 12). Vehicle mice were treated with vehicle (olive oil) for 6 weeks (n = 3) or 12 weeks (n = 3).

### 3.11. Human liver and pancreatic ductal adenocarcinoma (PDAC) specimen collection and institutional review board (IRB) approval

The human liver study included 8 patients diagnosed with liver diseases who underwent surgical resection. Liver tissue specimens were collected from both cirrhotic and normal liver tissue. Specimens were histologically confirmed by a pathologist.

The PDAC study included both treatment-naïve patients and those who had received neoadjuvant chemoradiotherapy followed by chemotherapy and radiation therapy, prior to surgical resection. Tumor specimens and non-tumor pancreatic tissue adjacent to the tumor (hereafter referred to as “adjacent non-tumor pancreas”) were collected during surgery and verified by a pathologist. Inclusion criteria required histopathological confirmation of PDAC.

All tissue collections were conducted in accordance with institutional review board (IRB) approval (DF/HCC 02-240), and written informed consent was obtained from each patient before surgery. The biopsied tissues were snap-frozen in O.C.T using liquid nitrogen and stored at –80°C until further processing.

### 3.12. Frozen tissue preparation for histology

Resected mouse organs, mouse colons prepared as “Swiss rolls”, or human tissue samples, were submerged in Tissue-Tek O.C.T. Compound (4583, Sakura) and snap-frozen in cold 2-methylbutane (MX0760-1, Supelco) in a beaker surrounded by dry ice under a fume hood. Frozen tissue was sectioned on a cryostat (CM1950, Leica) at 10 μm thickness and mounted onto positively charged glass slides. For Swiss rolls, tissue sections were examined under a microscope to ensure that the full intestinal architecture was preserved throughout the entire Swiss roll.

### 3.13. Masson’s trichrome staining for frozen tissue sections

Tissue sections were rehydrated in 4 °C PBS for 10 minutes, and a water-proof circle was drawn around the tissue using a hydrophobic barrier PAP pen (H-4000, Vector). Tissue sections were fixed in methanol for 4 minutes and washed in ultrapure water for 5 minutes. Sections were re-fixed in Bouin’s solution (15990, EMS) at 56 °C for 1 hour. Sections were rinsed in running tap water for 1 minute or until the yellow color disappeared. Sections were placed in Weigert’s hematoxylin (AB245882, Abcam) for 5 minutes and then rinsed in running tap water for 10 minutes. Sections were briefly rinsed in ultrapure water and then stained with Biebrich scarlet-acid fuchsin solution (26033-25, EMS) for 10 minutes. Sections were briefly rinsed in ultrapure water and then differentiated in a 2.5% phosphomolybdic-phosphotungstic acid solution for 15 minutes (19402-10 and 19502-5, EMS). Sections were transferred (without rinsing) to aniline blue solution (1.25 g of aniline blue was mixed with 1 mL glacial acetic acid and 50 mL ultrapure water) (02570-25, Polysciences). Sections were rinsed in ultrapure water and differentiated in 1% acetic acid for 3 minutes. Sections were briefly washed in ultrapure water, then in 95% ethanol for 20 seconds, followed by 100% ethanol for 20 seconds three times, and finally in xylene for 5 minutes twice. After the tissue was fully dry, mounting medium (SP15-100, Fisher) was added to the slides and covered with a coverslip. Slides were imaged the next day using a microscope (Nikon Instruments). Collagen quantification was performed on QuPath (version 0.6) by color deconvolution [40].

### 3.14. Picrosirius red staining for frozen tissue sections

Tissue sections were rehydrated in 4 °C PBS for 10 minutes and then fixed in methanol for 4 minutes and washed in ultrapure water for 5 minutes. Picro-Sirius Red solution was prepared by adding 0.5 grams of Sirius Red F3B (365548, Sigma) to 500 mL of saturated aqueous solution of picric acid (P6744, Sigma). Tissue sections were stained in Picro-Sirius Red solution for 1 hour to reach an equilibrium staining. Tissue sections were then rinsed in acidified water (0.5% glacial acetic acid) twice for 1 minute each and rinsed in 100% ethanol three times for 1 minute each. Sections were cleared in Xylene for 3 minutes. After the tissue was fully dry, mounting medium (SP15-100, Fisher) was added to the slides and covered with a coverslip. Slides were imaged the next day using a microscope (Nikon Instruments).

### 3.15. Hematoxylin and eosin (H&E) staining for tissue frozen sections

Tissue sections were removed from the freezer and allowed to warm to room temperature. Tissue sections were fixed in cold acetone at -20 °C for 10 minutes and subsequently washed in PBS three times for 5 minutes each. Tissue sections were stained in Gill 3 Hematoxylin (72611, Epredia) for 45 seconds and were rinsed in 3 changes of distilled water. Tissue sections were then stained in Eosin Y Alcoholic (6766007, Epredia) for 1 minute. Slides were differentiated in 50% ethanol for 1 minute, 75% ethanol for 1 minute, 95% ethanol for 1 minute, and 100% ethanol twice for 1 minute each. Sections were cleared in Xylene twice for 3 minutes each. After the tissue was fully dry, mounting medium (SP15-100, Fisher) was added to the slides and covered with a coverslip. Slides were imaged the next day using a microscope (Nikon Instruments).

### 3.16. Verhoeff Van Gieson elastin staining

Tissue sections were rehydrated in 4 °C PBS for 10 minutes and then fixed in methanol for 4 minutes and washed in ultrapure water for 5 minutes. A modified Verhoeff Van Gieson elastic stain kit (HT25A, Sigma) was used for staining elastin in the tissue sections. The instructions for the elastic stain kit were followed.

### 3.17. Image analysis of allysine staining

Image quality was further enhanced using artificial intelligence to denoise the raw data with the same image acquisition software (Nikon NIS-Elements). Digitized images were processed using ImageJ (Fiji distribution, version 1.54p). Non-specific staining and uneven fluorescence background were corrected using the rolling ball algorithm (radius: 50 pixels). Images were then converted to binary masks by applying the IsoData algorithm (the “Default” thresholding method in ImageJ), and total positive allysine staining area was quantified within regions of interest (ROI) in the binary masks.

To place ROIs for the distal part of the large intestine, the first 2 mm from the anus to the rectum was excluded, and then a freehand line was drawn to define a 14-mm line along the muscularis mucosa starting at the 2 mm borderline. An ROI was placed on the 14-mm-long colon, including the epithelial layer to the muscularis propria but excluding serosa.

To quantify the percentage of allysine-positive staining in the whole colon, an ROI was placed to cover the whole large intestine tissue section. The percentage of allysine-positive staining was calculated by dividing the area of positive staining by the total tissue area within the ROI.

To quantify the percentage of allysine-positive staining in other tissues, four separate ROIs were selected within the tissue sections, avoiding major vessels and ducts. The percentage was calculated by dividing the area of positive staining by the total tissue area within each ROI.

### 3.18. Statistics

Data are reported as the means ± SD. Differences between the two groups were tested with two-tailed unpaired and paired t tests. Differences between the two groups across multiple conditions were tested using multiple unpaired two-tailed t-tests, with p-values adjusted for multiple comparisons using the false discovery rate (FDR) method. Differences among more than two groups were tested with one-way analysis of variance (ANOVA), followed by Tukey’s post hoc test, with P < 0.05 considered significant. Agreement in allysine-positive area percentage between two adjacent sections was evaluated using Bland–Altman analysis. All statistical analyses were performed using GraphPad Prism 10.4 (GraphPad software).

## Acknowledgment

We thank Hua Ma for providing AL-NP standards for HPLC analysis, Iris Y. Zhou, and Yongtao Wang for providing CCl_4_ and CDAHFD liver tissue samples.

## Authors contributions

M.L.F. conceived the project. M.L.F. and P.C. supervised the study and secured the funding. D.L. performed tissue staining and analytical assays with support from P.P. D.L. prepared the DSS model. C.B. and P.P. sectioned the tissue blocks. I.C.H. performed SHG and assisted with imaging analysis. I.R.E. and K.K.T. collected human tissue samples. J.C.T.C. provided helpful discussions. J.T. reviewed all the histological staining. D.L. and M.L.F. generated the figures. D.L., M.L.F., and P.C. wrote the paper with the input from all authors. All authors reviewed and approved the final version of the paper.

## Supplementary Materials

Supplementary Fig. S1 to S5

## References

1. Rockey, D.C., P.D. Bell, and J.A. Hill, Fibrosis—a common pathway to organ injury and failure. New England Journal of Medicine, 2015. 372(12):p. 1138–1149.

2. Henderson, N.C., F. Rieder, and T.A. Wynn, Fibrosis: from mechanisms to medicines. Nature, 2020. 587(7835):p. 555–566.

3. Mutsaers, H.A., et al., The impact of fibrotic diseases on global mortality from 1990 to 2019. Journal of Translational Medicine, 2023. 21(1): p. 818.

4. Cox, T.R., The matrix in cancer. Nature Reviews Cancer, 2021. 21(4):p. 217–238.

5. Piersma, B., M.-K. Hayward, and V.M. Weaver, Fibrosis and cancer: A strained relationship. Biochimica et Biophysica Acta (BBA)-Reviews on Cancer, 2020. 1873(2): p. 188356.

6. Winkler, J., et al., Concepts of extracellular matrix remodelling in tumour progression and metastasis. Nature communications, 2020. 11(1): p. 5120.

7. Nissen, N.I., et al., The roles of collagens and fibroblasts in cancer, in Biochemistry of Collagens, Laminins and Elastin. 2024, Elsevier. p. 419–434.

8. Chen, W., et al., Lysyl oxidase (LOX) family members: rationale and their potential as therapeutic targets for liver fibrosis. Hepatology, 2020. 72(2):p. 729–741.

9. Rieder, F., et al., Fibrosis: cross-organ biology and pathways to development of innovative drugs. Nature Reviews Drug Discovery, 2025: p. 1–27.

10. Rønnow, S., et al., Elastin, in Biochemistry of Collagens, Laminins and Elastin. 2024, Elsevier. p. 279–289.

11. Singh, D., V. Rai, and D.K. Agrawal, Regulation of collagen I and collagen III in tissue injury and regeneration. Cardiology and cardiovascular medicine, 2023. 7(1): p. 5.

12. Yamauchi, M. and M. Sricholpech, Lysine post-translational modifications of collagen. Essays in biochemistry, 2012. 52: p. 113–133.

13. Kong, W., et al., Collagen crosslinking: effect on structure, mechanics and fibrosis progression. Biomedical Materials, 2021. 16(6): p. 062005.

14. Kisseleva, T. and D.A. Brenner, Mechanisms of fibrogenesis. Experimental biology and medicine, 2008. 233(2):p. 109–122.

15. Hedtke, T., et al., A comprehensive map of human elastin cross-linking during elastogenesis. The FEBS journal, 2019. 286(18):p. 3594–3610.

16. van der Slot-Verhoeven, A.J., et al., The type of collagen cross-link determines the reversibility of experimental skin fibrosis. Biochimica et Biophysica Acta (BBA)-Molecular Basis of Disease, 2005. 1740(1):p. 60–67.

17. Mäki, J.M., et al., Lysyl oxidase is essential for normal development and function of the respiratory system and for the integrity of elastic and collagen fibers in various tissues. The American journal of pathology, 2005. 167(4):p. 927–936.

18. Trackman, P.C. and M.V. Bais, Measurement of lysyl oxidase activity from small tissue samples and cell cultures, in Methods in cell biology. 2018, Elsevier. p. 147–156.

19. Palamakumbura, A.H. and P.C. Trackman, A fluorometric assay for detection of lysyl oxidase enzyme activity in biological samples. Analytical biochemistry, 2002. 300(2):p. 245–251.

20. Aslam, T., et al., Optical molecular imaging of lysyl oxidase activity–detection of active fibrogenesis in human lung tissue. Chemical Science, 2015. 6(8):p. 4946–4953.

21. Zhong, Y., et al., Lysyl oxidase regulation and protein aldehydes in the injured newborn lung. American Journal of Physiology-Lung Cellular and Molecular Physiology, 2022. 322(2):p. L204–L223.

22. Waghorn, P.A., et al., High sensitivity HPLC method for determination of the allysine concentration in tissue by use of a naphthol derivative. Journal of Chromatography B, 2017. 1064: p. 7–13.

23. Wang, H., et al., An in situ activity assay for lysyl oxidases. Communications biology, 2021. 4(1): p. 840.

24. Aronoff, M.R., et al., Imaging and targeting LOX-mediated tissue remodeling with a reactive collagen peptide. Nature Chemical Biology, 2021. 17(8):p. 865–871.

25. Carlson, J.C., et al., Superbright Bioorthogonal Turn-on Probes. Angewandte Chemie (International ed. in English), 2013. 52(27).

26. Ko₡lmel, D.K. and E.T. Kool, Oximes and hydrazones in bioconjugation: mechanism and catalysis. Chemical reviews, 2017. 117(15):p. 10358–10376.

27. Cordes, E. and W. Jencks, On the mechanism of Schiff base formation and hydrolysis. Journal of the American Chemical Society, 1962. 84(5):p. 832–837.

28. Wang, S., et al., Saline accelerates oxime reaction with aldehyde and keto substrates at physiological pH. Scientific Reports, 2018. 8(1): p. 2193.

29. Hussey, G.S., T.J. Keane, and S.F. Badylak, The extracellular matrix of the gastrointestinal tract: a regenerative medicine platform. Nature Reviews Gastroenterology & Hepatology, 2017. 14(9):p. 540–552.

30. Chen, X., et al., Second harmonic generation microscopy for quantitative analysis of collagen fibrillar structure. Nature protocols, 2012. 7(4):p. 654–669.

31. Bettenworth, D., et al., A global consensus on the definitions, diagnosis and management of fibrostenosing small bowel Crohn’s disease in clinical practice. Nature Reviews Gastroenterology & Hepatology, 2024. 21(8):p. 572–584.

32. Rieder, F., et al., Fibrosis in IBD: from pathogenesis to therapeutic targets. Gut, 2024. 73(5):p. 854–866.

33. Suzuki, K., et al., Analysis of intestinal fibrosis in chronic colitis in mice induced by dextran sulfate sodium. Pathology international, 2011. 61(4):p. 228–238.

34. Melgar, S., A. Karlsson, and E. Michaëlsson, Acute colitis induced by dextran sulfate sodium progresses to chronicity in C57BL/6 but not in BALB/c mice: correlation between symptoms and inflammation. American Journal of Physiology-Gastrointestinal and Liver Physiology, 2005. 288(6):p. G1328–G1338.

35. Hammerich, L. and F. Tacke, Hepatic inflammatory responses in liver fibrosis. Nature Reviews Gastroenterology & Hepatology, 2023. 20(10):p. 633–646.

36. Ning, Y., et al., Molecular MRI quantification of extracellular aldehyde pairs for early detection of liver fibrogenesis and response to treatment. Science translational medicine, 2022. 14(663):p. eabq6297.

37. Waghorn, P.A., et al., Molecular magnetic resonance imaging of lung fibrogenesis with an oxyamine-based probe. Angewandte Chemie International Edition, 2017. 56(33):p. 9825–9828.

38. Ning, Y., E.A. Akam-Baxter, and P. Caravan, Extracellular Aldehyde Sensing Probes for In Vivo Imaging. Accounts of Chemical Research, 2025. 58(14):p. 2203–2215.

39. Wahsner, J., et al., 68Ga-NODAGA-indole: an allysine-reactive positron emission tomography probe for molecular imaging of pulmonary fibrogenesis. Journal of the American Chemical Society, 2019. 141(14):p. 5593–5596.

40. Ruifrok, A., Quantification of histochemical staining by color deconvolution. Analytical and quantitative cytology and histology/the International Academy of Cytology [and] American Society of Cytology, 2001.

