## Supplementary Figures for "Rapid imaging of lysyl oxidase activity and fibrogenesis with a turn-on fluorophore"

### Supplementary Fig. 1

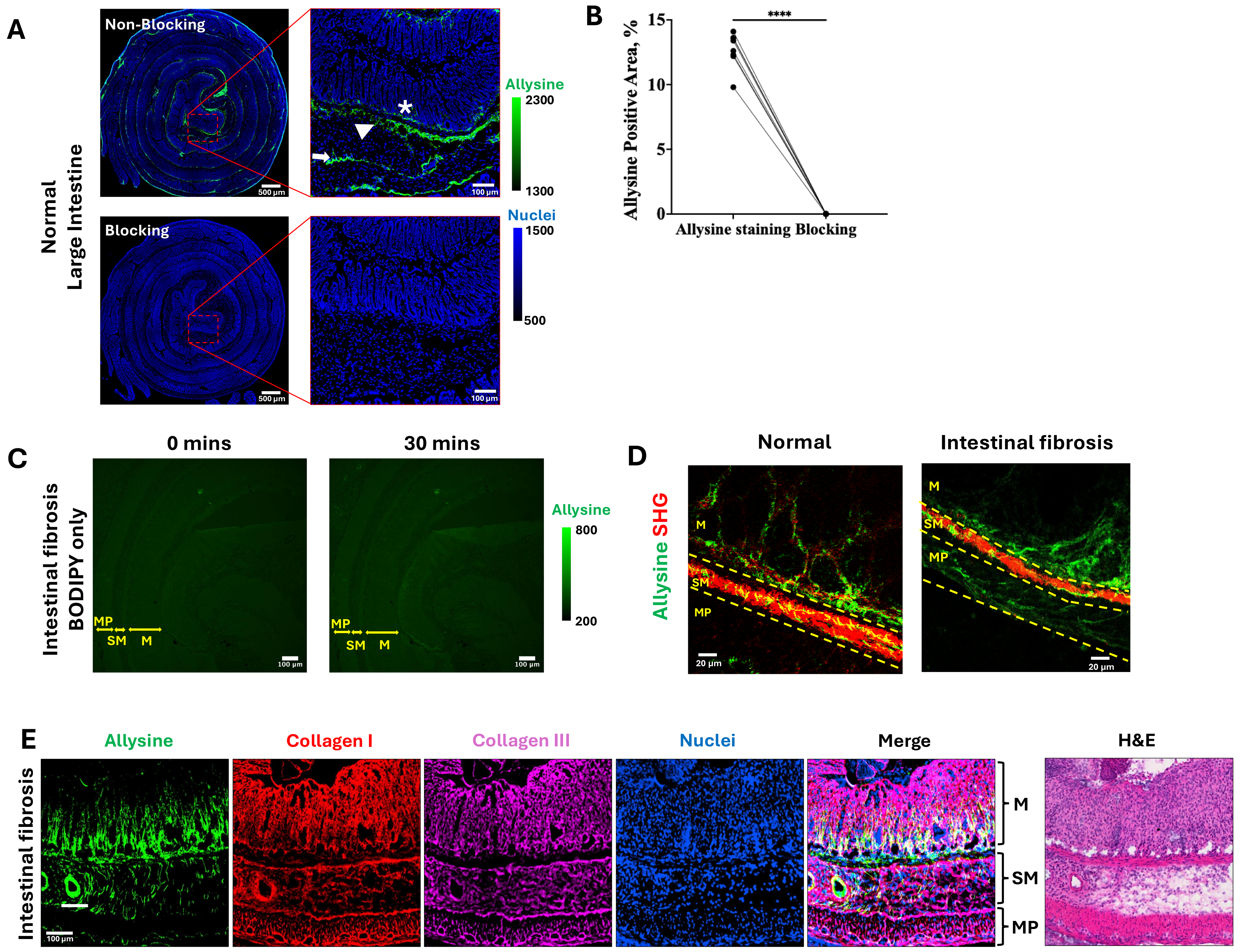


**Supplementary Fig. 1. Staining method validation. (A)** The large intestine of 9-week-old naïve male C57BL/6 mice was stained by allysine staining (n = 8) (Top). A higher magnification image showed the allysine distribution in the distal part of the large intestine. The asterisk highlights the allysine in the mucosa layer. The arrowhead points to the allysine in the submucosa layer. The arrow points to the allysine in the layer between the circular and longitudinal muscle. When blocking the large intestine with methoxyamine hydrochloride, no positive staining was observed. **(B)** Quantification of allysine-positive area percent in the distal part of the intestine (n = 8). Blocking the allysine in the tissue with methoxyamine hydrochloride eliminated all the allysine-positive staining (n = 8). ****$P < 0.0001$, paired t test, two-tailed. **(C)** Real-time allysine imaging of fibrotic tissue of the large intestines (n = 3). Tissues were not pre-tagged with BCN-PEG3-Oxyamine. Tissue sections were imaged immediately after the addition of methyltetrazine-BODIPY. The first image was acquired right after adding methyltetrazine-BODIPY (less than 1 min). During the 30-minute incubation, BODIPY was not activated, and the fluorescent signal remained largely unchanged. **(D)** Two-photon images of the middle part of the large intestines (n = 3) stained by allysine staining (green), and collagen was visualized by SHG. **(E)** Allysine staining and immunofluorescence staining of collagen I and collagen III were performed on the same fibrotic colon tissue section. The same tissue sections were stained by hematoxylin and eosin (H&E) staining. M: mucosa; SM: submucosa; MP: muscularis propria.

#### Supplementary Fig. 2

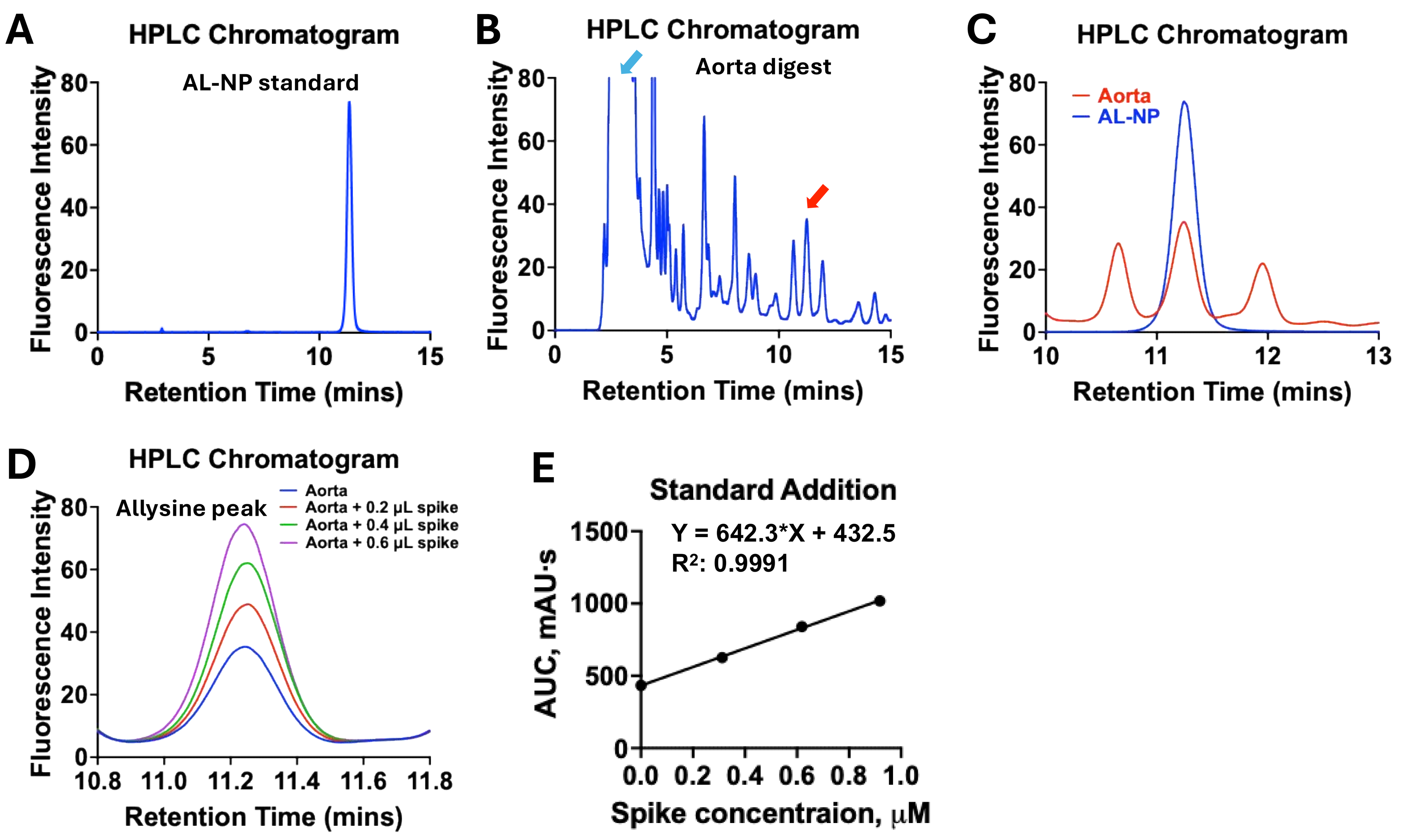


**Supplementary Fig. 2. Allysine quantification assay by high-performance liquid chromatography (HPLC). (A)** HPLC chromatogram of allysine-naphthol (AL-NP) standard. **(B)** HPLC chromatogram of aorta labeled with 2-naphthol-6-sulfonate. The blue arrow indicates the excess 2-naphthol-6-sulfonate peak. The red arrow indicates the AL-NP peak. **(C)** HPLC chromatogram of aorta labeled with 2-naphthol-6-sulfonate overlaid with AL-NP standard. **(D)** HPLC chromatogram of aorta spiked with AL-NP standard using the standard addition method. **(E)** Allysine concentration determination in the aorta sample by standard addition. The absolute value of the x-intercept of the linear regression is the allysine concentration in the injected HPLC sample.

#### Supplementary Fig. 3

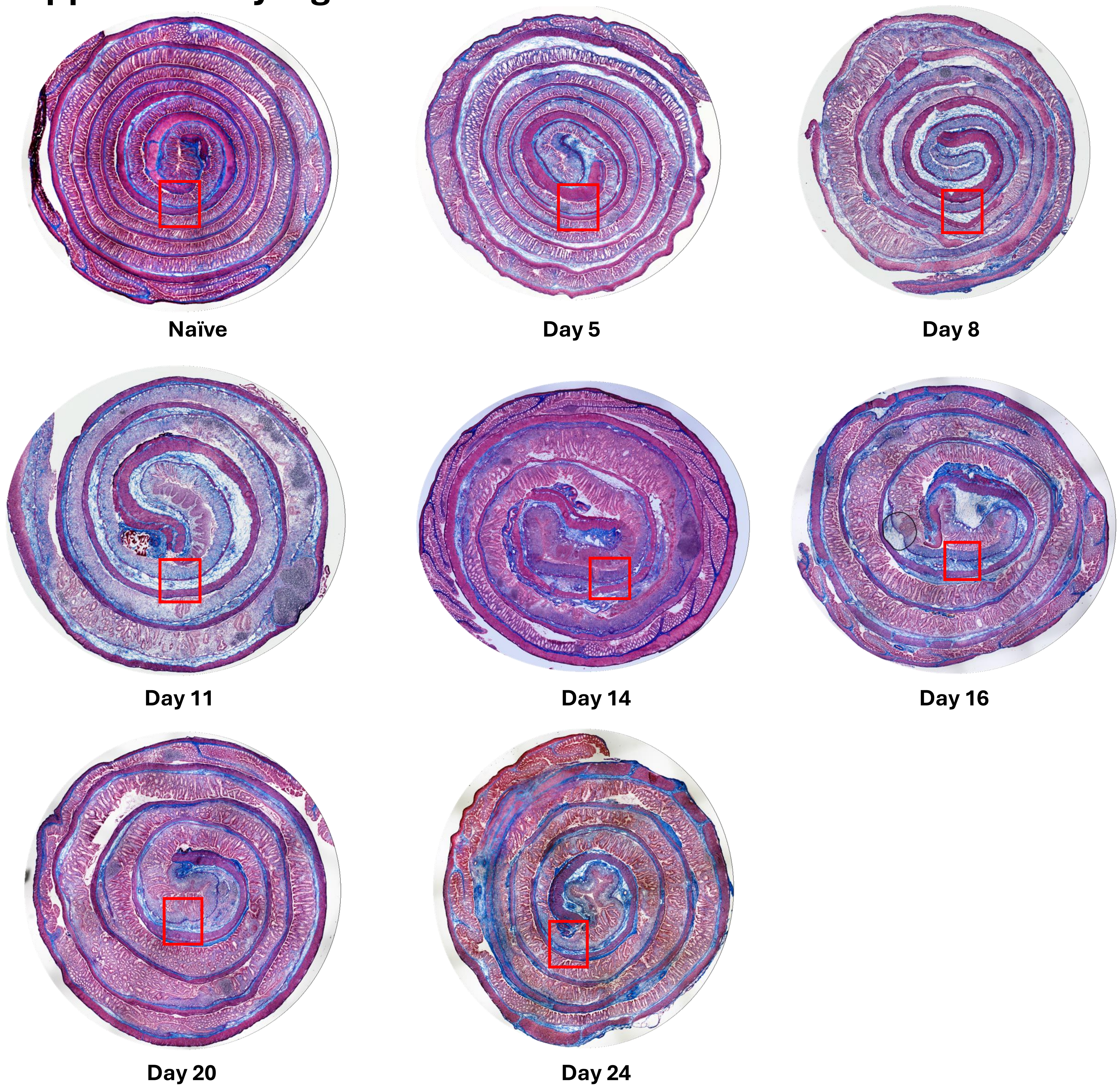


**Supplementary Fig. 3. Representative Masson's Trichrome staining of the entire large intestine for the natural history of the dextran sodium sulfate (DSS)-induced intestinal fibrosis mouse model.** The whole large intestine was stained by Masson's Trichrome staining in naïve mice (n = 12), and on days 5 (n = 4), 8 (n = 4), 11 (n = 4), 14 (n = 8), 16 (n = 3), 20 (n = 3) and 24 (n = 3) post-DSS induction. The red boxes delineate where the high magnification images are used in Figure 3A.

### Supplementary Fig. 4

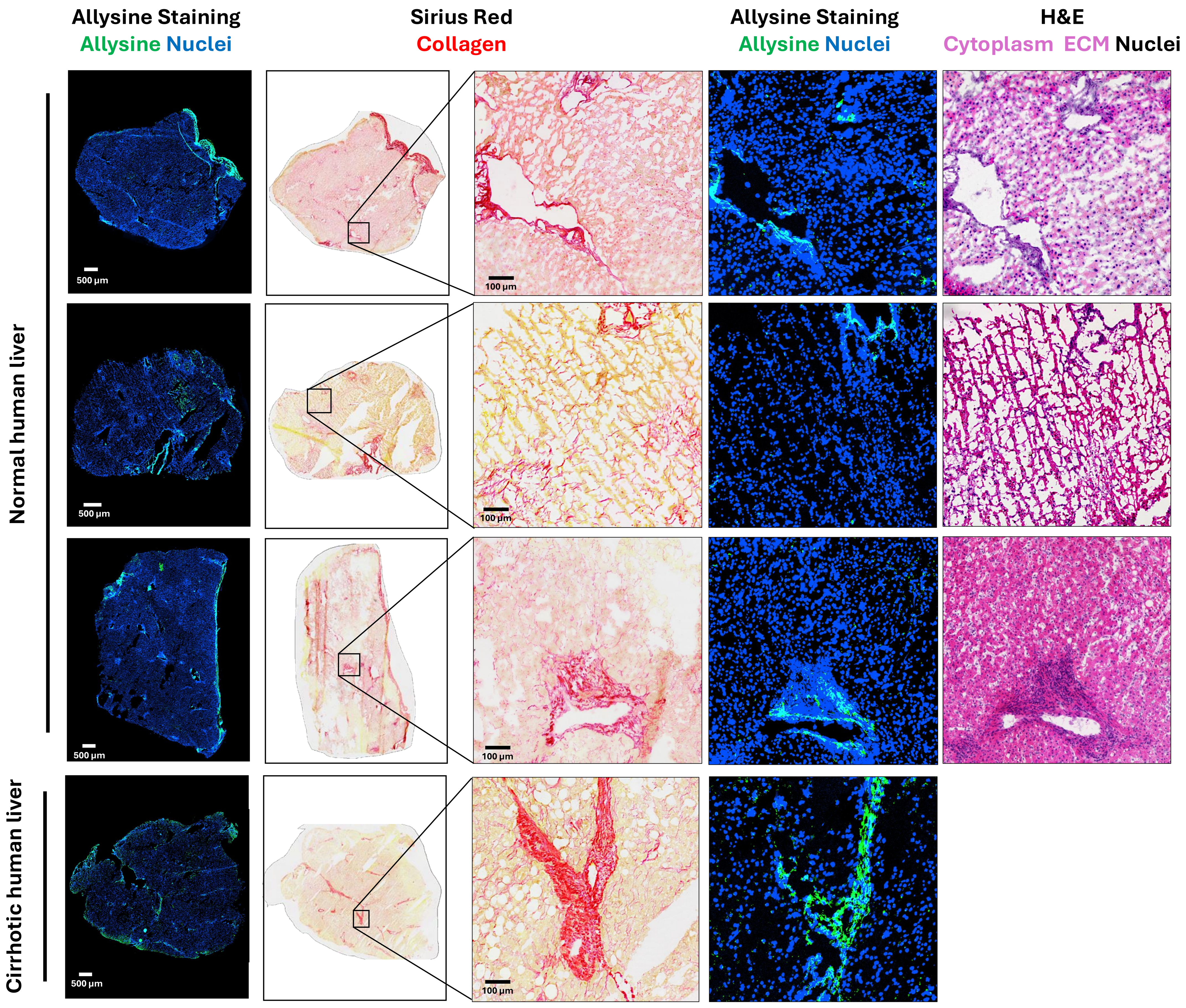


**Supplementary Fig. 4. Cirrhotic and normal human liver specimens from patients with liver diseases.** The allysine and collagen levels were mapped in the entire liver tissue section by allysine and Sirius red staining in adjacent tissue sections. Hematoxylin and eosin staining visualized the tissue morphology.

### Supplementary Fig. 5

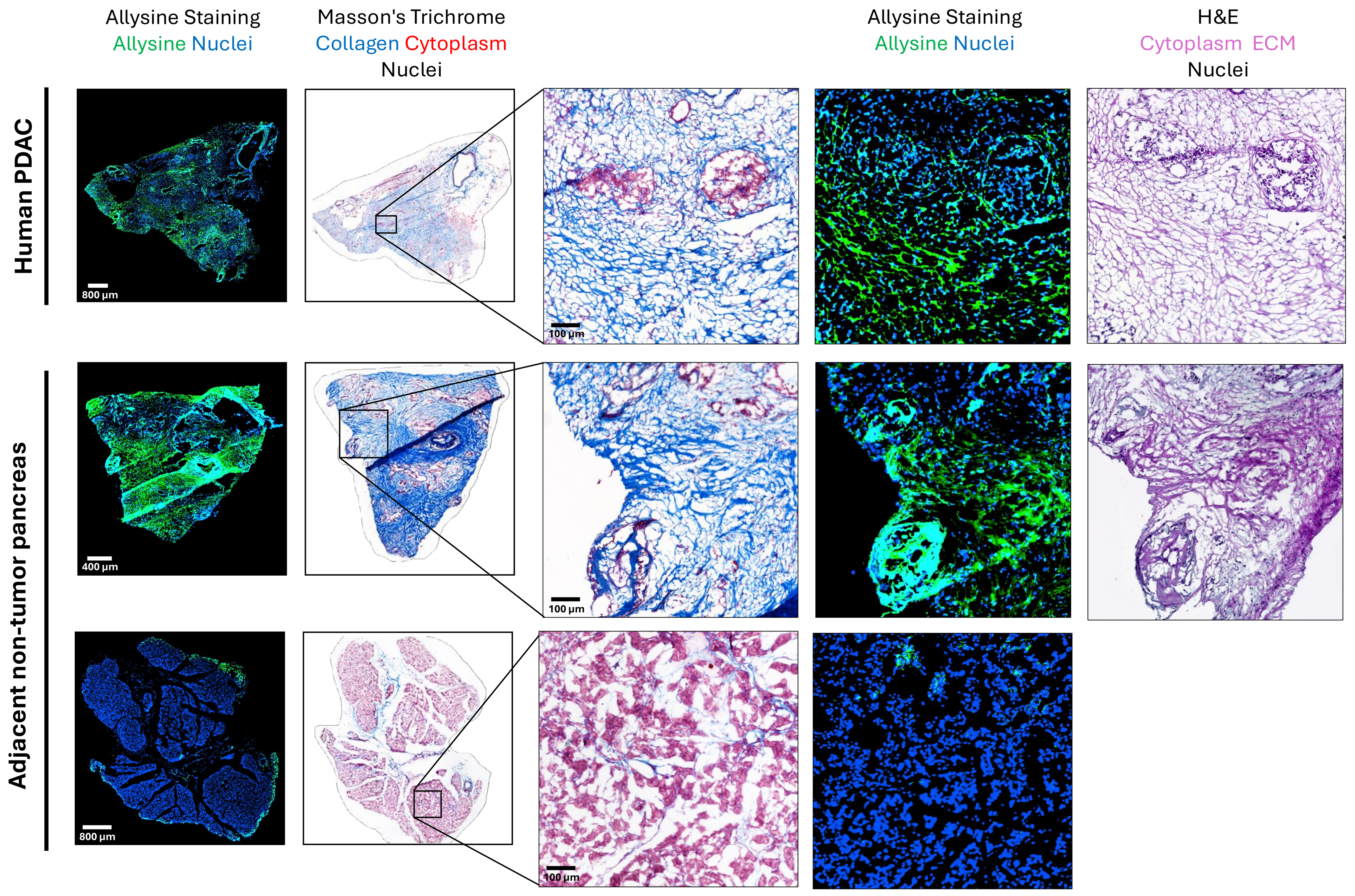


**Supplementary Fig. 5. Fibrotic human pancreatic ductal adenocarcinoma (PDAC).** Human pancreatic ductal adenocarcinoma (PDAC) and adjacent non-tumor pancreas specimens were stained using allysine, Masson's Trichrome and H&E stains. The allysine and collagen levels were mapped in the entire pancreas tissue section by allysine and Masson's Trichrome staining. H&E staining showed the tissue morphology. The distribution of allysine is consistent with the collagen-rich TME. Allysine staining only maps active fibrosis in TME.
